# The Age-Dependent Decline in Neuron Growth Potential in the CNS is Associated with an Age-Related Dysfunction of Neuronal Mitochondria

**DOI:** 10.1101/2021.05.14.444223

**Authors:** Theresa C. Sutherland, Arthur Sefiani, Darijana Horvat, Taylor E. Huntington, Yuanjiu Lei, A. Phillip West, Cédric G. Geoffroy

## Abstract

The age of incidence of Spinal Cord Injury (SCI) and the average age of people living with SCI is continuously increasing. In contrast, SCI is extensively modelled in young adult animals, hampering translation of research to clinical application. While there has been significant progress in manipulating axon growth after injury, how it is impacted by aging impacts this is still unknown. Aging is associated with a decline in mitochondrial functions, whereas mitochondria are essential to successful neurite and axon growth. Using isolation and culture of adult cortical neurons, we have analyzed mitochondrial changes in 2-, 6-, 12- and 18-month mice. We observed reduced neurite growth in older neurons. Older neurons also showed dysfunctional respiration, reduced membrane potential, and altered mitochondrial membrane transport proteins; however mitochondrial DNA (mtDNA) abundance and cellular ATP were increased. Taken together, these data suggest dysfunctional mitochondria in older neurons are involved in the age-dependent reduction in neuron growth. Both normal aging and traumatic injury are associated with mitochondrial dysfunction, posing a challenge for an aging SCI population as the two elements can compound one another to worsen injury outcomes. The results of this study highlight this as an area of great interest in CNS trauma.

## 1. Introduction

Spinal Cord Injury (SCI) is the second most common cause of paralysis and results in varying degrees of motor and sensory dysfunction over the lifetime of the patient. The international incidence rate of SCI is approximately 13 cases per 100,000 population [1], while the incidence in the United States is approximately 54 cases per million population, and 17,730 new cases each year [2, 3]. The demographic of the SCI population has seen an important shift in recent decades. The average age at injury is increasing, as is the age of the SCI population in general. The current average age of incidence in the US is 43 years old [3]. Within this population there are two peaks in incidence, one in young (20-30 years) and one in aging (≥65 years) adults [1]. As a result of the increase of SCI in middle-aged and aging populations and the fact that people with SCI live longer, the average of people who reported paralysis due to a SCI is today ~48 [4]. There has been a lack of consideration of the effects of aging on SCI, and especially axon growth, in current SCI research. In contrast to the human SCI population the majority of pre-clinical research is performed in young adult rodents (2-4 months), and less than 0.35% of experimental rodents used are 12 months or older, which represents 40 years of age in humans [5]. This is likely to be a significant impediment to translating pre-clinical research to viable clinical therapies.

Significantly, an age-dependent decline in axon growth has been reported in a variety of model organisms, both invertebrates and vertebrates. This decline is influenced by both extra-neuronal [6] and neuron intrinsic factors [7]. It has been previously reported that even genetic manipulation of neuron-intrinsic factors aimed at promoting axon growth are sensitive to age and decrease in efficiency and efficacy at various ages [8]. This observation suggests that there are other significant factors at play in the age-dependent decline in axon growth potential, and other elements that can be targeted to promote axon growth in aging neurons. One factor of particular interest is neuronal mitochondria. There is extensive evidence that functional decline of mitochondria contributes to normal aging throughout the body, including prominent roles in age-related processes such as cellular senescence, chronic inflammation, and decline in stem cell activity [9].

The mitochondrial, or free-radical, theory is a prominent theory of aging, which places mitochondrial decline as a fundamental mechanism of biological and cellular aging [9–12]. This theory has been proposed for decades with reactive oxygen species (ROS) at its core, however, in recent years evidence has highlighted other facets of mitochondrial dysfunction to be important in the process of cellular aging. Extensive work still implicates mitochondria as a central figure, but also leaves many questions still to be answered [12]. Mitochondria are involved in a plethora of vital cellular functions including energy production, intracellular homeostasis, calcium balance, and the metabolism of various dietary substrates, and immunity [10]. It is established that mitochondrial function throughout the body is reduced with age, and this decline has been associated with a variety of diseases, including age-related neurodegeneration [10]. Mitochondrial dysfunction is considered one of the hallmarks of the aging brain [13].

The most prominent role for mitochondria is cellular respiration and energy production, which is significant in SCI as neuron and axon growth requires significant cellular energy. Any dysfunction in OXPHOS and the electron transport chain (ETC) may have a significant impact on the ability of aging neurons to support axon sprouting and growth. Additionally, these organelles are also involved in calcium (Ca^2+^) homeostasis and buffering, vital to neuronal function. Mitochondria at the synapse are involved in buffering Ca^2+^ to regulate neurotransmission [14]. The mitochondrial permeability transition pore (mPTP) becomes more active with increased ROS and oxidative stress, and prolonged opening of this channel effects both ROS balance and Ca2+ homeostasis. Other mitochondrial membrane channels, such as translocase of outer membrane (TOM) [15, 16] and inner membrane (TIM) [17, 18] complexes, are essential to the translocation of a variety of bio-active proteins and molecules into the mitochondria [19–21]. OXPHOS Complex I subunit protein GRIM19 [22, 23] plays a role in maintaining mitochondrial membrane potential (ΔΨM) [22] and also in transport across the membrane [19]. Alteration in expression and function of these proteins and complexes, as well as decreases in membrane potential, affect mitochondrial ability to effectively import essential molecules as well as maintain cellular Ca^2+^ levels.

Mitochondria play important roles in normal aging [9], in axon growth [24–27], and in the progression of SCI [28–30]. Significantly, mitochondrial dysfunction and detrimental oxidative stress is associated with both normal aging as well as central nervous system (CNS) trauma. This poses a challenge for an aging SCI population that falls at the intersection of these two factors as they may compound each other and worsen the outcome for aging SCI patients. Using a new system of adult cortical neuron isolation and culture, we compared neuronal mitochondrial functions between four ages of mice-2-months (young), 6-months (young adult), 12-months (middle-aged) and 18-months (old). We observed a variety of detrimental changes in neuronal mitochondria in the aging CNS that will have a significant effect on both neuronal health and the ability to promote neurite growth after injury. Collectively, our results suggest for the first time an important role for mitochondria functions in the age-dependent decline in axon growth potential. This also may suggest the promotion of neuronal mitochondrial function as a key element to promote axon growth in injured neurons regardless of age or time post injury. These observations need to be expanded *in vivo* and strategies to manipulate the mitochondria analyzed in detail.

## 2. Materials and Methods

### Animals

This study uses both male and female wild-type C57Bl/6 mice of 4 age groups; 2-months, 6-months, 12-months and 18-months. All procedures were conducted according to Institutional approved Animal Use Protocols.

### Isolation of Adult Cortical Neurons

Adult cortical neurons were isolated in an enriched population from the cortex using the Mitlenyi MACS system- the Adult Brain Dissociation Kit (130-107-677) and Adult Neuron Isolation Kit (130-126-603). The mice were euthanized using CO^2^ and the brains removed. The cortex was isolated using blunt dissection under a microscope and transferred to a MACS C-tube. Samples were dissociated with two mice per tube. This was then dissociated using the appropriate pre-set protocol on the Miltenyi gentleMACS octo-dissociator with heaters. The manufacturer’s recommended protocols were followed for both the Adult Brain Dissociation Kit and Adult Neuron Isolation Kit (using LS-columns) with minor adjustments. During this dissociation and isolation protocol the neurons are stripped of their neurites and axons, allowing this to be a model for axonal injury and neurite growth *In Vitro*. This protocol was used as the basis for all of the subsequent analysis performed in this study.

### RT-qPCR for Isolated Cell Population Purity

RNA was extracted from both the negative fraction (neurons) and positive fraction using Directzol RNA micro-prep columns (Zymo, R2061) directly following MACS dissociation and isolation (D0), as well as after 2- and 4-days *In Vitro* and Seahorse analysis. RNA concentration was measured using NanoDrop (Thermo Fisher) and samples were calculated accordingly. cDNA was synthesised using Quantabio cDNA Synthesis kit (Quanta, 95047), followed by qPCR with Quantabio Perfecta Sybr Fastmix + Rox (Quanta, 95073) run on ViiA7 Real Time PCR system (Life Technologies), as previously published [31]. Primers were used to identify the main cellular constituents of the isolated populations: Neuron- MAP2 (F: 5’-CTG GAG GTG GTA ATG TGA AGA TTG; R: 5’-TCT CAG CCC CGT GAT CTA CC-3’) and NeuN (F: 5’-AAC CAG CAA CTC CAC CCT TC-3’; R: 5’-CGA ATT GCC CGA ACA TTT GC-3’); Astrocytes- GFAP (F: 5’-CTA ACG ACT ATC GCC GCC AA-3’; R: 5’-CAG GAA TGG TGA TGC GGT TT-3’) and Glast (F; 5’-CAA CGA AAC ACT TCT GGG CG-3’; R: 5’-CCA GAG GCG CAT ACC ACA TT-3’); and Oligodendrocytes- Oligo2 (F; 5’- GAA CCC CGA AAG GTG TGG AT-3’; R: 5’-TTC CGA ATG TGA ATT AGA TTT GAG G-3’); as well as β-actin (F: 5’-CTC TGG CTC CTA GCA CCA TGA AGA-3’; R: 5’-GTA AAA CGC AGC TCA GTA ACA GTC CG-3’) for normalization. This analysis was performed in triplicate on available young neuron samples on DO (2-6-month) (N=2), and in young (2-6-month) (N=2) and aging (12-18-month) (N=2) samples collected after seahorse analysis.The neuron enrichment in the negative fraction was calculated as -ΔCT of NeuN against GFAP and Glast using the formula: *-ΔCT = -(ΔCT NeuN – (SQRT(ΔCT GFAP^2^ + ΔCT Glast^2^)))*.

### Analysis of Neurite Growth In Vitro

The enriched cortical neuron population resulting from MACS dissociation and isolation of female mice was plated onto poly-D-lysine (PDL) and laminin coated #1.5 thickness glass culture plates. At the time of plating, these cells were without axons and significant neurites due to the dissociation process. These cells were cultured in MACS Neuro media (Miltenyi) with penicillin (50units/ml)/streptomycin (50μg/ml) (Corning), L-Alanyl-L-glutamine (Sigma-Aldrich, 2mM), B-27 Plus Supplement (Gibco, 1X) and brain-derived neurotrophic factor (BDNF; Tonbo Biosciences, 1μg/ml), with 50% media replacement every second day, and fixed in 8% paraformaldehyde (PFA; 15min) at the end of 7 days. Immunocytochemistry was performed on PFA fixed cells for βIII-Tubulin (1:500, BioLegend) and cells were imaged using the ImageXpress automated confocal system (Molecular Devices) using 10x objective (3 samples/age). Neurite growth was analyzed from the 10x βIII-Tubulin images using NeuronJ (ImageJ) to measure average neurite length/cell, total neurite length/cell, the number and length of primary (1°), secondary (2°) and tertiary (3°) neurites, and the number of branching points visible (N=50-75 cells/age). Cells were counted as neurons and measured only if they were βIII-Tubulin positive and the neurites were ≥10μm.

### qPCR of Mitochondrial DNA

After MACS isolation (as described previously) the enriched cortical neuron cells were immediately disrupted and lysed for RNA and DNA extraction using an AllPrep DNA/RNA Micro kit (Qiagen, 80284). The protocol was performed to the manufacturer’s recommendation for this kit. Mitochondrial DNA analysis by qPCR was performed on the extracted DNA as previously published [32]. Briefly, DNA was subjected to qPCR using PerfeCTa SYBR Green FastMix (Quanta) and the CFX384 Real-Time Syctem (BIO-RAD). Each sample analysed consisted of the complete cortex from 2 mice pooled, and was run in triplicate normalized against nuclear-encoded 18s using the 2^-ΔΔCt^ method. Relative expression (%) analysis was performed as described using primers specific to multi-copy cytosolic ribosomal DNA sequence 18s (F: 5’-CTT AGA GGG ACA AGC GGC G-3’; R: 5’-ACG CTG AGC CAG TCA GTG TA-3’) and mitochondrial ribosomal DNA sequence 16s (F: 5’-GTT ACC CTA GGG ATA ACA GCG C-3’; R: 5’-GAT CCA ACA TCG AGG TCG TAA ACC-3’). Only two genes were used here due to the small amount of DNA in each sample. These were analysed as 2-, 6-, 12-, and 18-month samples (N=2/age, in triplicate), and as grouped young (2- and 6-) and older (12- and 18-) (N=4/group, in triplicate).

### Analysis of Mitochondrial Respiration

Cortical neurons were isolated from female mice as described above, with each sample consisting of the cortices from 2 mice pooled together. Isolated cortical neurons were plated at 40,000 cells/well onto a 96 well Seahorse XFe96 plates treated with PDL and laminin and left to adhere and begin neurite sprouting over 4 days. After 4 days *in vitro* these cells were analyzed on the Agilent Seahorse XFe Analyzer using the Mito Stress Test kit. This kit uses three drug cocktails added into the assay media over time to alter the cell respiration-Oligomycin (2.5μM), Trifluoromethoxy carbonylcyanide phenylhydrazone (FCCP; (2.0μM) and Rotenone/Antimycin A (0.5μM). The oxygen consumption rate (OCR) is measured to assess mitochondrial respiration, and the extracellular acidification rate (ECAR) and proton efflux rate to assess glycolysis. This analysis was run to the manufacturer’s suggested protocol. Due to small cell yield and low baseline OCR the 2- and 6-month neurons, and the 12- and 18-month neurons, were grouped into ‘young’ and ‘older’ respectively for analysis to increase the number of wells/group (N=4/group).

### Western Blot for OXPHOS protein complexes

After isolation of an enriched cortical neuron population (from 2 brains/sample, female) using the previously described protocol, the isolated cell populations were immediately placed in 20μl RIPA protein extraction buffer (EMD Millipore) with HALT Protease/Phosphatase inhibitors (ThermoFisher Scientific) and PMSF (Cell Signalling Technology). Protein extraction was performed on ice for 30min with intermittent mixing. The protein supernatant was separated by centrifuging at 3000xg for 30min and stored at −80 until use. Sample concentrations were measured using NanoDrop (Thermo Fisher) and 50μg of protein was run for each sample (N=3/ age). Protein was run on Novex 10% Tris-Glycine Mini Gels (Thermo Fisher) using Novex Tris-Glycine SDS Running Buffer (Thermo Fisher) and NuPAGE Sample Reducing Agent (Thermo Fisher). Gels were run for 2 hours and 10 min at 80V and transferred using the Trans-Blot SD Semi-Dry Transfer cell system (Bio-Rad). After transfer the LI-COR Revert 700 Total Protein Stain (LI-COR) was applied, per manufacturer’s protocol, and imaged using LI-COR Odyssey CLx Imaging System (LI-COR). Following this the membranes were treated using SuperSignal Western Blot Enhancer (Thermo Fisher) for 10 min before blocking in 5% Milk in TBST for 1 hour. After washing in 0.05% TBST Total OXPHOS Rodent Antibody Cocktail (Abcam; 1:2000) was added to the membranes and incubated overnight at room temperature. After washing off the primary antibody, the secondary antibody (Anti-Mouse IgG HRP-linked; Cell Signaling; 1:2000) was added and incubated for 1 hour, followed by Supersignal West Femto Maximum Sensitivity Chemiluminescent Substrate (Thermo Fisher) for 5 min. The samples were analysed 6 times in repeated western blots. The blots were imaged using the LI-COR Odyssey Fc Imaging System (LI-COR Biosciences), chemiluminescent setting. Protein expression, as measured by band intensity (mean greyscale value; MGV), was normalized to the total protein stain.

### Functional Assay for Mitochondrial ATP Production

Neurons were isolated from female mice as described above with MACS. Cells were manually counted under a microscope using 0.4% Trypan Blue (VWR, 97063-702) and inserted into a Levy Counting Chamber (Hausser Scientific™ 3900) to determine cell number in each sample. Each sample consisted of the cortical neurons preparation from 2 mice pooled (N=3 samples/age) and each sample was run in triplicate as described below. The enriched cortical neuron population was immediately placed in 50μl RIPA protein extraction buffer (EMD Millipore) with HALT Protease/Phosphatase inhibitors (ThermoFisherScientific) and PMSF (Cell Signalling Technology) to lyse the cells and mitigate ATPase activity. After a 30 min incubation at 4 °C with intermittent mixing, the supernatant was separated by centrifuging at 20,000 xg for 20 min. Immediately following lysing, the relative ATP concentration of isolated neurons was measured using the bioluminescence-based ATP Determination Kit (Invitrogen, A22066) following the manufacturer’s recommended protocol. Briefly, 6 μL of each sample, standard, or control was mixed with 60 μL of Standard Reaction Buffer (1 mM dithiothreitol, 0.5 mM D-luciferin, 1.25 μg/mL firefly luciferase, Component E Reaction Buffer) before being loaded onto a 384-well polystyrene μCLEAR® bottom plate (Greiner, 781091) and incubated at 28°C for 10 min. Relative Light Units (RLU) were measured at 28°C using the Synergy 2 SL Microplate Reader with Gen5™ software (BioTek).

Known ATP standards were run alongside samples to determine ATP concentration. The ATP concentration of lysed from 2-, 6-, 12-, and 18-month old cortical neurons was determined using a linear equation developed from the ATP standards: *Concentration of ATP (nM) = 0.0035 x Relative Light Units (RLU)* (R^2^ = 0.9753). Using the volume of the assay, number of cells lysed, and ATP concentration of each sample, the approximate number of ATP molecules in cortical neurons from each age group was determined. The number of ATP/cell was calculated and expressed as a percentage change relative to the youngest group in the analysis (2-month, or 2-6-months), this measure is shown in the resulting graphs.

### Flow Cytometry Analysis of Membrane Potential

JC-1 (5,5,6,6’-tetrachloro-1,1’,3,3’ tetraethylbenzimi-dazoylcarbocyanine iodide) is a lipophilic cationic dye frequently used to analyses mitochondrial membrane potential (ΔΨM). JC-1 has normally green fluorescence until it forms into aggregates, which have red fluorescence. In healthy cells with a normal ΔΨM, JC-1 enters and accumulates in the mitochondria at high levels, and forms red fluorescent (PE) aggregates. In unhealthy or stressed cells, whose mitochondria have increased membrane permeability and loss of electrochemical potential, JC-1 enters the mitochondria at decreased levels that are not sufficient for the formation of aggregates maintaining its normal green fluorescence (FITC) [33].

Cortical Neurons were isolated from female mice using the MACS protocol previously described, with one mouse/sample (N=8/age). Samples were isolated and analyzed two at a time young versus aged, 2-against 12-month and 6-against 18-months. Following isolation the samples were split in half, one half was treated with FCCP at 40μM for 5 minute as a control, before JC-1 was added to all the cell samples and incubated in Neuron media (37°C, 5% CO_2_) for 20 minutes. After this incubation, additional media was added to these samples and they were centrifuged. The cell pellet was suspended in FACS buffer (PBS, 5% FBS, 1% BSA, 0.05% Sodium Azide) and analyzed using a BD Fortessa flow cytometer with BD Diva software using (using PE, FITC and BV421 lasers). Styox Blue dead cell stain was added to the samples immediately prior to analysis. 50,000 events were analyzed per sample. These samples were first gated using forward (FCS) and side scatter (SSC), followed by height/area for single cells, and then live cells based on Sytox staining. From this population of live, single cells the Duel positive population (both PE and FITC) was analyzed for PE and FITC fluorescent intensity (MFI). This was expressed and analyzed as a red:green ratio (PE/FITC).

### Mitochondrial Membrane-associated Protein Immunocytochemistry

The cortical neurons plated and cultured for 7-days that were used for neurite analysis were also stained for either TOM20 (1:200, Proteintech), TIM23 (1:200, Santa Cruz Biotech) or GRIM19 (1:200, Santa Cruz Biotech). Cells were imaged using the 20x objective on a Zeiss Axio Observer system. The mean greyscale value (MGV) per area of each individual neuron was analyzed using ImageJ (N= 50-75/age).

### Data Analysis

Seahorse data was analysed in Agilent Wave and Seahorse Analytics XF software. Flow cytometry data was analysed using FlowJo (v10.7.1). Normally distributed data was analyzed using ANOVA with Tukey’s post-hoc test, and unpaired t-tests using Graphpad Prism. All histograms are representative of the data mean with error bars showing the standard error of the mean (SEM).

## 3. Results

### 3.1. Cell Population Purity of Enriched Cortical Neurons after Isolation and Culture

RT-qPCR was used to analyze the cellular constituents of the enriched cortical neuron population. Two neuron-associate genes, two astrocyte-associated genes and and oligodendrocyte-associated gene were analyzed, normalized against internal β-actin. Neuron enrichment was calculated as *-ΔCT = -(ΔCT NeuN – (SQRT(ΔCT GFAP^2^ + ΔCT Glast^2^)))*. RNA extracted from the neuron enriched negative fraction immediately following MACS dissociation and isolation (D0) showed a neuron enrichment -ΔCT of 11.65. After 2-days *In Vitro* with neuron specific culture methods the neuron-enriched population had a -ΔCT of 11.05, and RNA extracted after 4-days in culture and analysis using the Agilent Seahorse showed a -ΔCT of 9.08. Olig2 expression was not observed in samples from 2-days and 4-days *in vitro*. Following from this we are confident the data presented below are from a highly enriched neuronal population.

### 3.2. Neurite growth potential declines in mature cortical neurons In Vitro with age

Analysis of *In Vitro* cortical neuron neurite outgrowth cultured for 7DIV from 2-, 6-, 12- and 18-month old mice showed decreases with age as well as altered patterns of growth (Figure 1). The total length of neurite growth per cell decreased with age, though this was only significant (P<0.05) between the 2-month and both the 12- and 18-month neurons (Figure 1A). The significance of this decrease is magnified (P<0.0005) when comparing grouped young (2- and 6-month) and older (12- and 18-month) neurons (Figure 1Ai). The average neurite length also significantly decreases in older neurons compared to young both between the four separate ages (Figure 1B) and in grouped ages (Figure 1Bi). The number of primary neurites was similar in young and older neurons (Figure 1C and Ci), however the length of the longest primary neurite decreased significantly between young and older neurons (Figure 1D and Di). Conversely, the number of secondary neurites was increased in older neurons compared to young (P<0.05) and the number of branching points (between primary, secondary and tertiary neurites) is also significantly increased in older neurons (Figure 1E and Ei). This suggests not only a decrease in neurite growth potential with age but also increased branching in aged neurons compared to their younger counterparts (Figure F-Gi).

**Figure 1.**
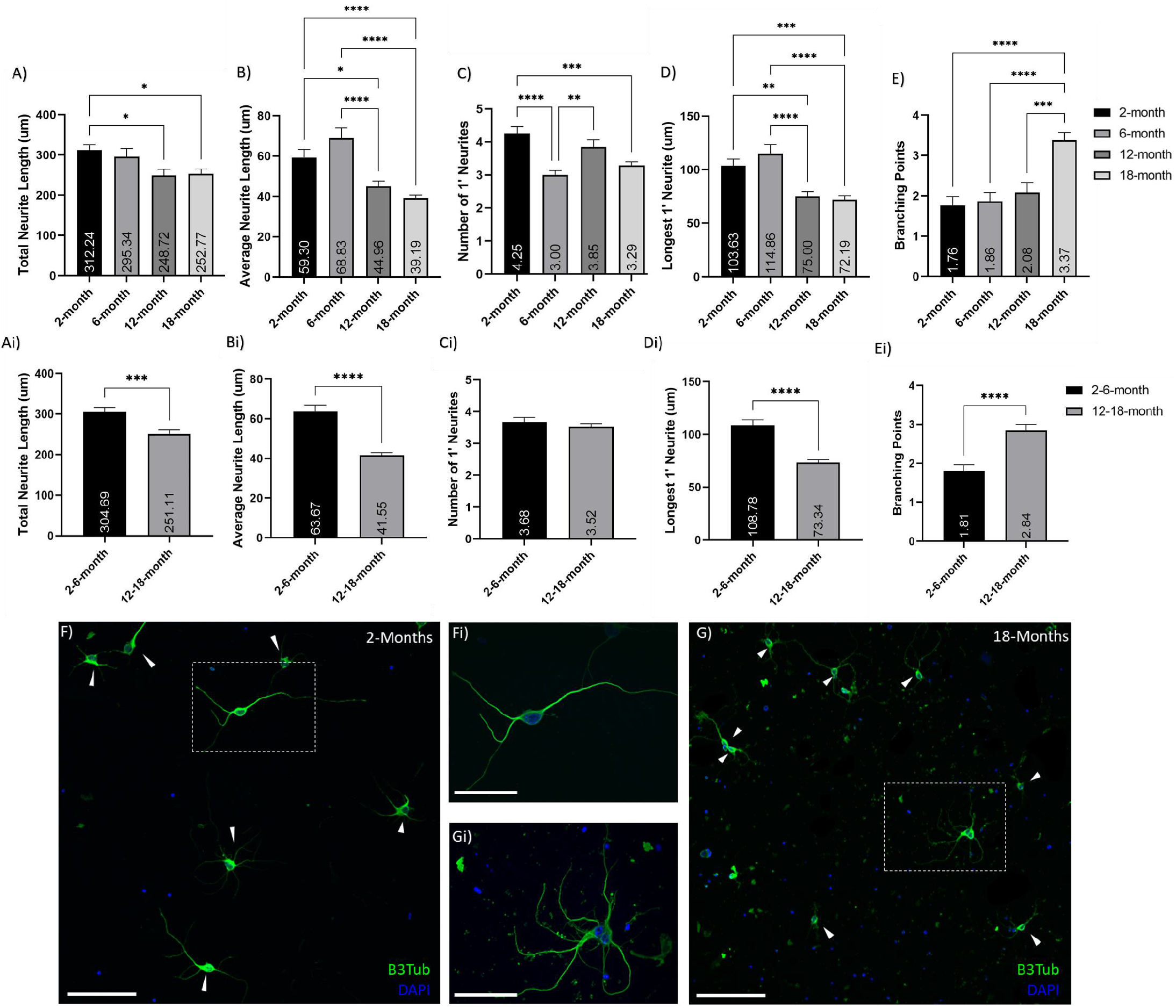
Reduction of neurite outgrowth with age. Histograms of neurite analysis and feature quantifications comparing between A-E) 2-, 6-, 12- and 18-month old neurons; as well as A-E) grouped young (2- and 6-month) and older (12- and 18-months) neurons.These show decreased neurite growth and increases branching in older neurons compared to young. A) and Ai) show age-related decreases in total neurite growth per cell. B) and Bi) show decreases in the average neurite length with age. C) and Ci) show the number of primary (1’) neurites per cell. D) and Di) show a decline in primary neurite length with age. E) and Ei) indicate an increase in neurite branching with age. This pattern is illustrated in the representatice 20x confocol images of F) 2-month and G) 18-month cortical neuron cultures after 7-days *in vitro;* stained with βIII-Tubulin (Green) and DAPI (Blue). Inset Fi) and Gi) show 60x magnification of neurons. These images also show the increased debris and less healthy look of the old neurons after 7 DIV. *\* (P<0.05), ** (P<0.005), *** (P<0.0005), **** (P<0.0001). N= 50-75 cells/age. Graphs show mean and SEM. Arrows indicate individual neurons. Scale bars are 100μm for F and G; 50 μm for Fi and Gi*.

### 3.3. Expression of Mitochondrial DNA and mitochondrial DNA copy number changes with age

Analysis of mitochondrial copy number was performed using quantitative PCR of 18s (nuclear) and 16s (mitochondrial) genes. Due to the small sample size from isolated cortical neuron populations only one nuclear and one mitochondrial gene was used. Relative expression of the mitochondrial 16s ribosomal DNA sequence normalized to the nuclear-encoded 18s ribosomal DNA sequence was calculated using the 2-month samples as a baseline. Separately, both the 12- and 18-month neurons had significantly higher relative expression (%) than the 6-month neurons (P<0.005), and the 18-month were also significantly increased compared to the 2-month neurons (P<0.05) (Figure 2A). When grouped into young (2- and 6-month) and older (12-18-months) this relative expression was significantly increased in older neurons compared to their younger counterparts (t-test; P=0.0004)(Figure 2B). This indicated an increase in mtDNA with age.

**Figure 2.**
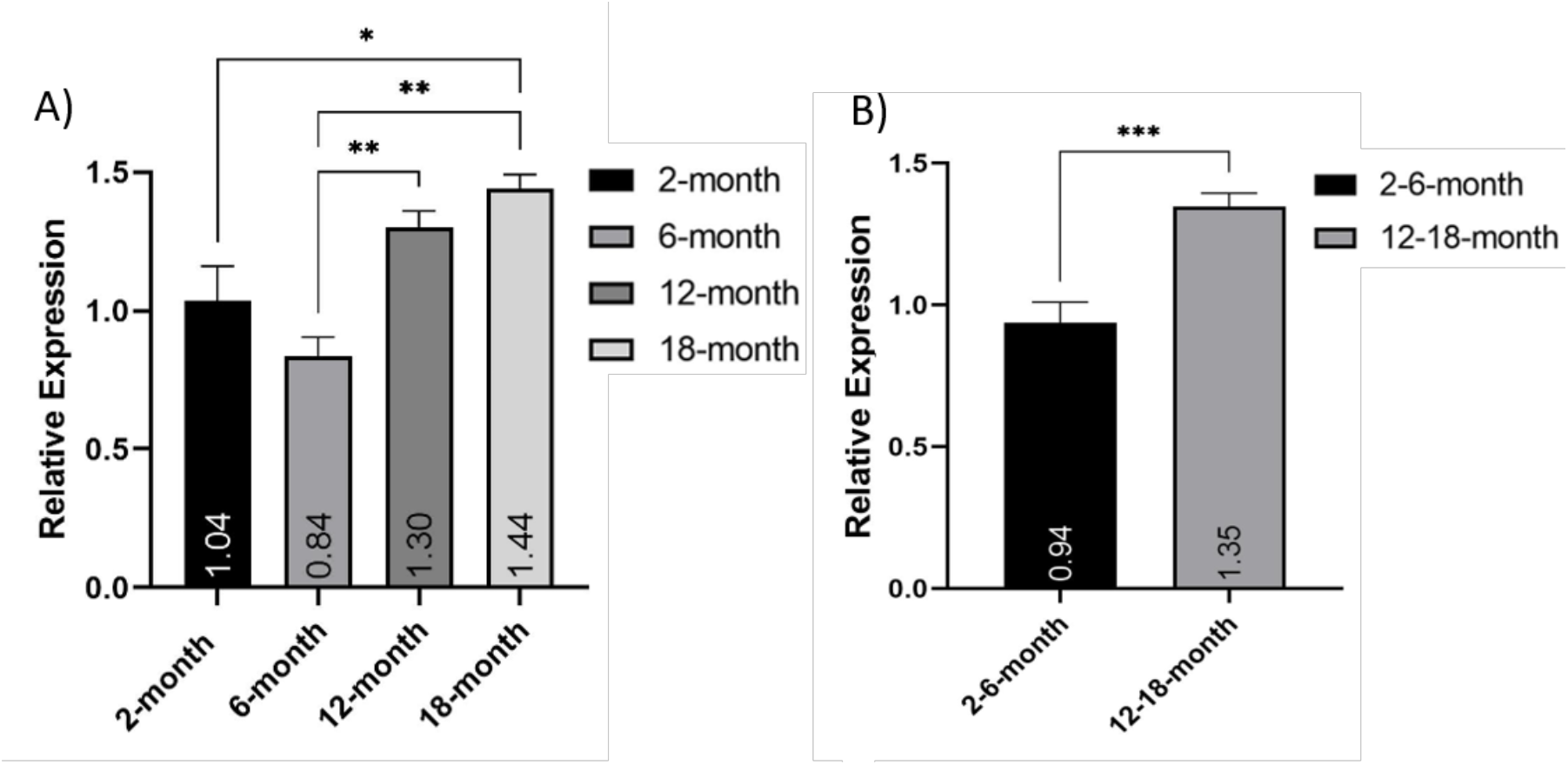
Increase in neuronal mitochondrial DNA expression with age. Relative expression of the mitochondrial *16s* ribosomal DNA sequence normalized to the nuclear-encoded *18s* ribosomal DNA sequence. A) comparing 2-, 6-, 12- and 18-month old isolated neuron populations (*N= 2);* and B) comparing grouped young (2- and 6-month) and older (12- and 18-month) (*N= 4). * (P<0.05), ** (P<0.005), *** (P<0.0005). Graphs show mean and SEM*.

### 3.4. Mitochondrial respiration of cultured mature cortical neurons changes with age

The Mito Stress Test (Agilent) measures both OCR, and ECAR at baseline levels and in response to drugs modulating mitochondria activity. First Oligomycin acts on Complex V to inhibit conversion of ADP to ATP; then FCCP alters membrane potential and stimulates maximal capacity of the respiratory chain; and finally, Rotenone inhibits Complex I and Antimycin A inhibits Complex III to suppress the ETC function (Figure 3A). This results in a response curve as seen in the 2-6-month neurons (Figure 3B). Interestingly we did not observe this strong response in the 12-18-month neurons, with only very subtle increases and decreases in OCR at the response time-points (Figure 3B). The baseline OCR for the 12-18-month neurons was significantly lower than that seen in the 2-6-month (Figure 3B), however there was no significance in the majority of parameters. From the OCR measurements, we observed a trend for decreased basal respiration (Figure 3C), maximal respiration (Figure 3D), ATP production (Figure 3E), proton leak (Figure 3F), and coupling efficiency (Figure 3G) in the 12-18-month group compared to the 2-6month. The non-mitochondrial oxygen consumption (Figure 3H) of older neurons showed a trend for increase compared to the younger counterparts. ECAR is associated with lactate efflux, and increases in this measure indicate cellular activation and proliferation. The ECAR measure for both 2-6-month and 12-18-month neurons remained fairly steady over the time course of this assay, and showed an increase in young neurons compared to older (Figure 3I). Plotting the OCR vs. ECAR, the young neurons show a tendency towards a more energetic metabolism compared to their more quiescent older counterparts. Together these measures suggest increased efficiency and respiratory capacity in younger cortical neurons, as well as increased ability to respond to respiratory stimulus, while older neurons are more quiescent and unresponsive. However, these isolated neurons measured quite low in this assay, especially the older groups, which calls for further exploration to strengthen these observations.

**Figure 3.**
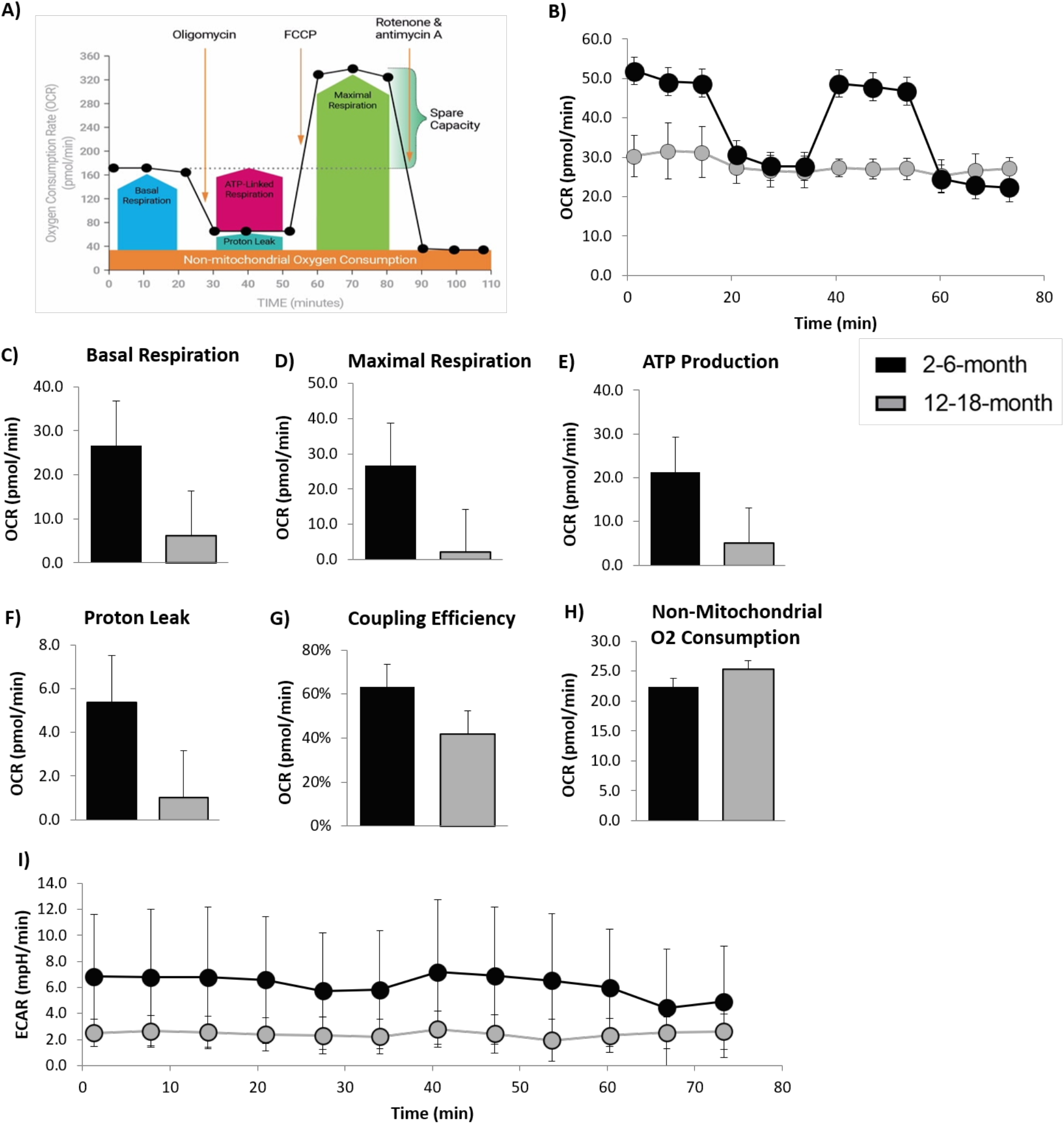
Alteration to Mitochondrial respiration in older neurons. A) Schematic of the injected drugs and time-course of the Seahorse Mito Stress Test (Agilent) showing the expected response curve and the respiratory measures calculated. B) Oxygen consumption rate (OCR) over the timecourse of the assay shows the expected response curve and higher OCR in the young (2- and 6-month) neurons compared to lower OCR and almost no responsiveness to the mitochoindrial active drugs in the grouped 12- and 18-month neurons (older). C) Basal respiration, D) Maximal respiration, E) ATP production, F) Proton leakage, and G) Coupling efficiency (%) were all increased in young cortical neurons compared to the older; while H) Non-mitochondrial oxygen consumption was decreased. I) The extracellular acidification rate (ECAR) over the time-course of the assay showed a similarly consistent response curve between young and older neurons, with the older neurons at a decreased level. There was no statistical significance to these trends as there was high wel to well variability. N=4/age. *Graphs show mean and SEM*.

### 3.5. Expression of mitochondrial OXPHOS complexes in the electron transport chain is consistent in cortical neurons of different ages

Expression of the five OXPHOS complexes of the ETC (C I – C V) was analyzed using western blot. Comparison of 2-, 6-, 12- and 18-month neuronal protein extract (directly after brain dissociation and cell separation) showed no significant alterations in expression of any of the five complexes (Figure 4A). When grouped into young (2-6-month) and older (12-18-month) no significant changes in any of the five complexes were observed (Figure 4B). Across all age groups Complex I and Complex V showed the highest expression. There was no statistical significance in these findings using either ANOVA or t-tests.

**Figure 4.**
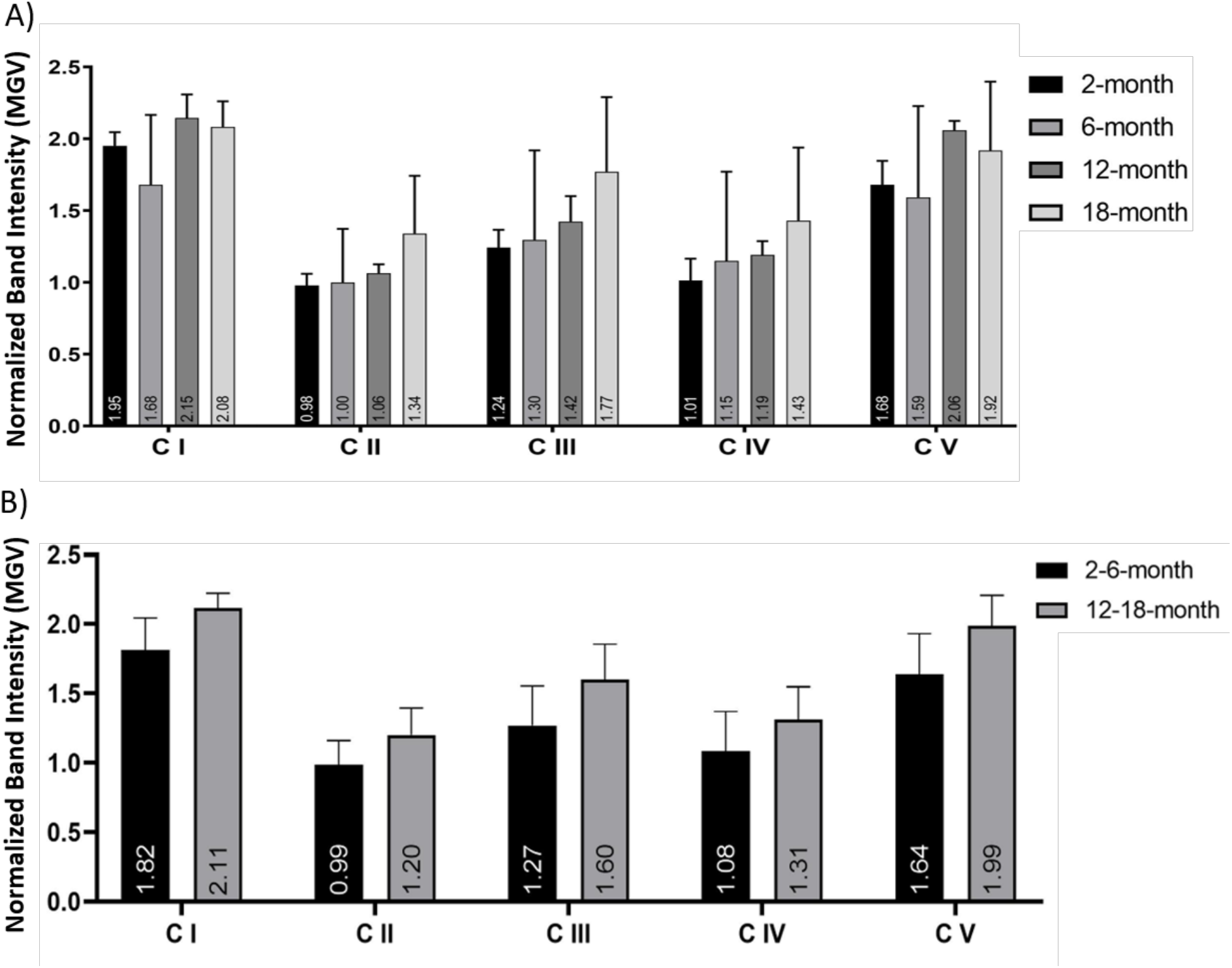
No significant change in the Mitochondrial OXPHOS Complex protein expression in cortical neurons. Histograms of the western blot band intensity, calculated as mean greyscale value (MGV) and normalized to the Total Protein stain, of proteins of all five OXPHOS Complexes (C I - C V) in the ETC comparing A) 2-, 6-, 12- and 18-month cortical neurons, and B) grouped young (2-6-month) versus aging (12-18-month) neurons, indicate no significant change in expression with age. This was not statistically signifcant in any comparison. *N= 3 samples/age, repeated 6 times. Histograms show mean and SEM*.

### 3.6. Intracellular ATP increases in neurons with age

Mitochondrial ATP production was measured using a bioluminescence-based ATP assay and compared between different age cohorts. Comparison of the number of intracellular ATP molecules in 2-, 6-, 12-, and 18-month old neurons showed a trend of increase with age, which was significant between the 2- and 12-month neurons (Figure 5A; P<0.05). When grouped into young (2- and 6-month) and older (12- and 18-month) neurons there was a significant increase in the relative number of intracellular ATP molecules (Figure 5B; P<0.05). There was an upward trend in the number of intracellular ATP molecules in respect to age and a simple linear regression equation was calculated to predict the number of intracellular ATP molecules based on the age of the cohort (P=0.1302, R^2^=0.7566)(Figure 5C).

**Figure 5.**
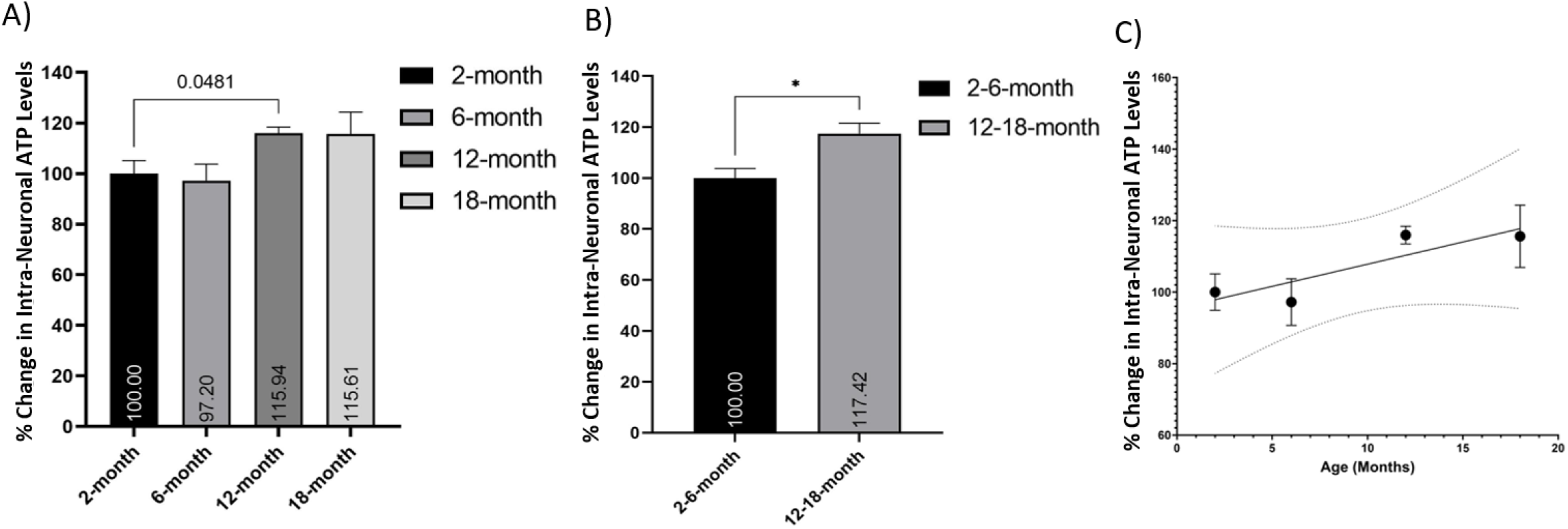
Increased retention of ATP molecules in older neurons. Histograms of the Number of Intracellular ATP Molecules per cell as measured using a bioluminescence-based ATP assay, expressed as a percentage change relative to the youngest group (Young or 2-months, respectively) comparing A) 2-, 6-, 12- and 18-month cortical neurons, and B) young (2-6-month) versus older (12- 18-month) neurons, indicate an trend of increase in intracellular ATP with age. C) This trend of increases can be expressed as a simple, non-significant linear relationship (P=0.1302, R^2^=0.7566) with 95% CI (dotted line). *\* (P<0.05). N= 3 samples/age, analysed in triplicate. Histograms show mean and SEM*.

### 3.7. The mitochondrial membrane potential of isolated cortical neurons changes with age

We next assess the change in mitochondrial membrane potential (ΔΨM) in freshly isolated cortical neurons of different ages. We used JC-1 flow cytometry to measure the PE:FITC mean fluorescent intensity (MFI) ratio in cortical neurons. Our data show that ΔΨM is altered in an age-dependent manner. Indeed, the The PE:FITC MFI ratio declined with age, however this is not significant in either grouped data or between the individual ages, due to high standard deviations in the data (Figure 6). This age-dependent alteration may have significant implications for mitochondrial function in cortical neurons.

**Figure 6.**
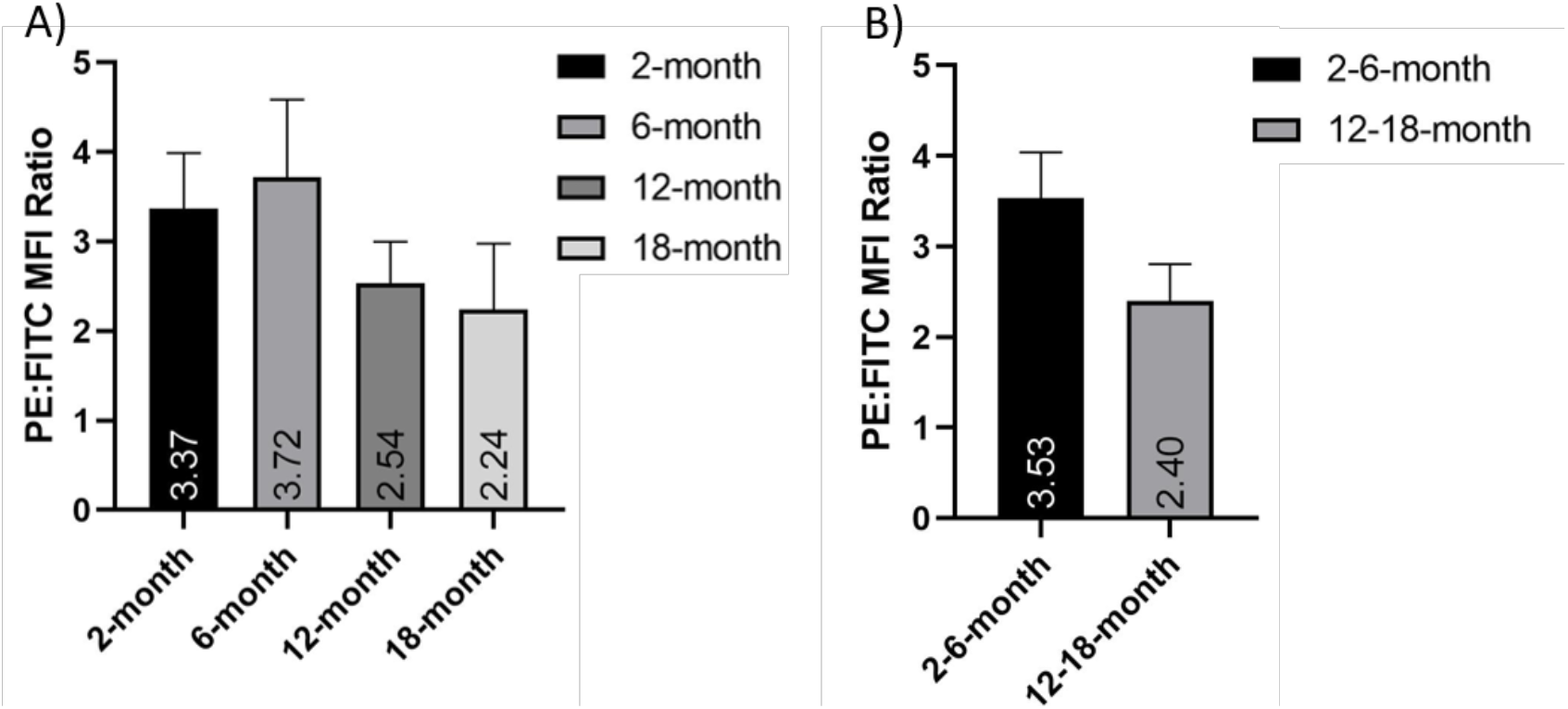
Depolarization of the mitochondrial membrane in older neurons. Histograms of the PE:FITC mean fluorescent intensity (MFI) ratio of JC-1 stained cortical neurons normalized to FCCP treated controls; both in A) 2-, 6-, 12- and 18-month old isolated neuron populations and B) grouped young (2- and 6-month) and older (12- and 18-month) neurons. These show a non-significant age-dependent decrease in mitochondrial membrane potential(*P=0.09 in B*). *N= 8. Graphs show mean and SEM*.

### 3.8. Mitochondrial membrane-associated proteins are altered in aging neurons

Immunocytochemistry for several mitochondrial membrane transport proteins showed a change in their expression with age (Figure 7). TOM20 is a prominent translocase of the outer mitochondrial membrane [15, 16] and was observed to decrease with age between the young (2- and 6-month) neurons and their older counterparts (12- and 18-months) (Figure 7A and Ai). It was also noted that the TOM20 immuno-staining appeared less visible in the long neurites of the older neurons, though the cell bodies were just as bright as the young (Figure 8). TIM23 is a translocase of the inner membrane that shuttles molecules into the mitochondrial matrix from the intermembrane space [17, 18]. TIM23 expression was surprisingly increased in older neurons compared to younger. Both 12- and 18-month neurons showed significantly increased TIM23 MGV/cell compared to the 2-month neurons (P<0.05 and P<0.0001 respectively). The 18-month neurons were also significantly higher in TIM23 than the 6-month (P<0.005) (Figure 7B). When grouped into young (2- and 6-month) and older (12- and 18-) the significance between young and old increased (P<0.0001) with the older neurons showing nearly double the TIM23 MGV/cell (Figure 7Bi). GRIM19 is a subunit of Complex I of the ETC [22, 23] that is involved in transport across the membrane [19] and maintenance of membrane potential [22]. GRIM19 expression showed a trend to decrease with age, though this was not significant except between the 6- and 18-month neurons (Figure 7C). When combined into young (2-6mnth) and older (12-18mnth) groups there was a significant decrease in GRIM19 expression (P<0.05) seen with age (Figure 7Ci).

**Figure 7.**
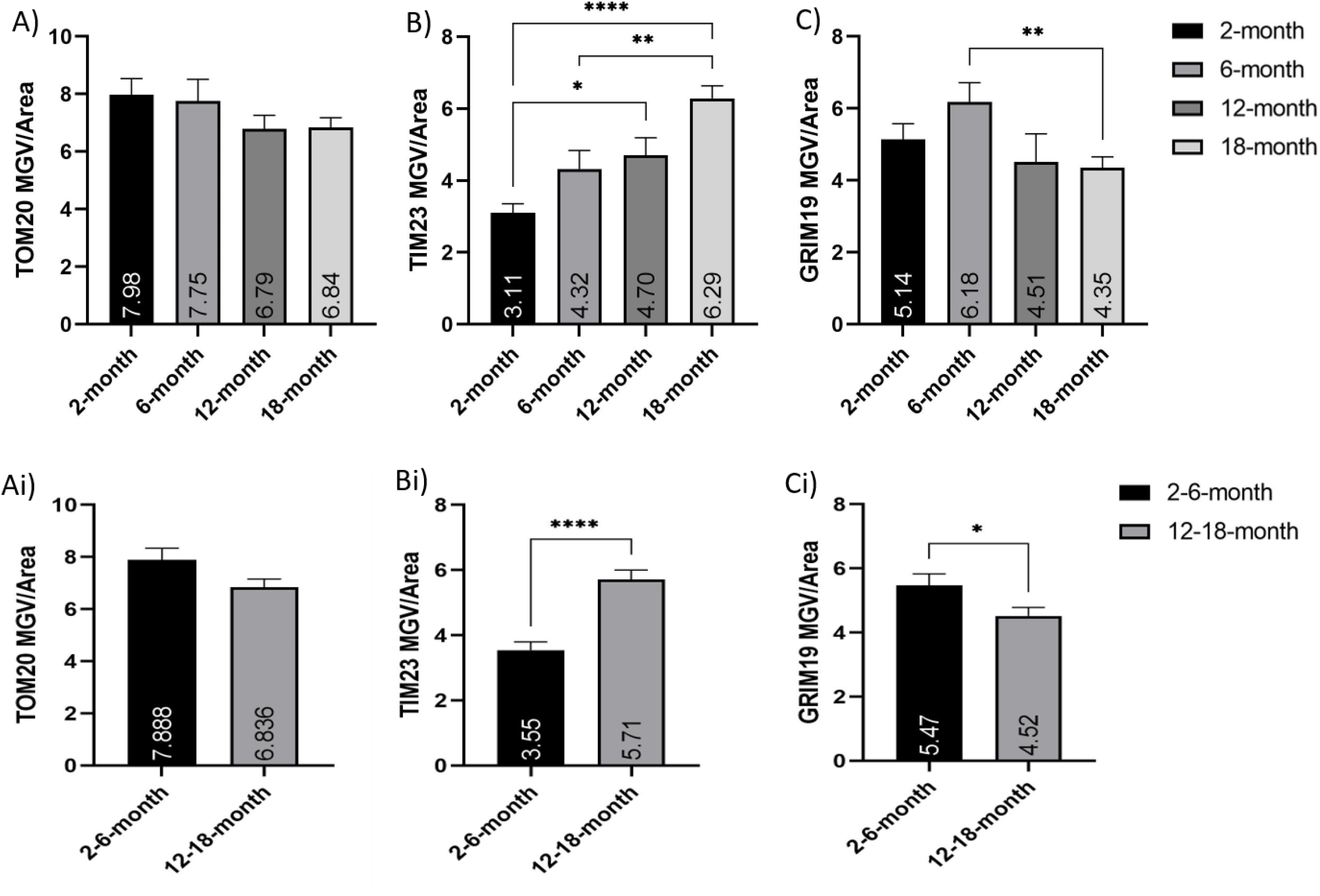
Alterations in expression of membrane transport associated proteins with age. Histograms of TOM20, TIM23 and GRIM19 fluorescent mean grayscale value/cell (MGV/cell) comparing A-C) grouped young (2- and 6-month) and older (12- and 18-months) neurons, as well as A-C) between 2-, 6-, 12- and 18-month old neurons. A) and Ai) show age-related a trend for a decrease in TOM20 expression per cell, though these were not significant (*P=0.1*). B) and Bi) show significant increases in the TIM23 expression with age. C) and Ci) show decreases in GRIM19 MGV per cell. *\* (P<0.05), ** (P<0.005), **** (P<0.0001). N= 50-75 cells/age. Graphs show mean and SEM*.

**Figure 8.**
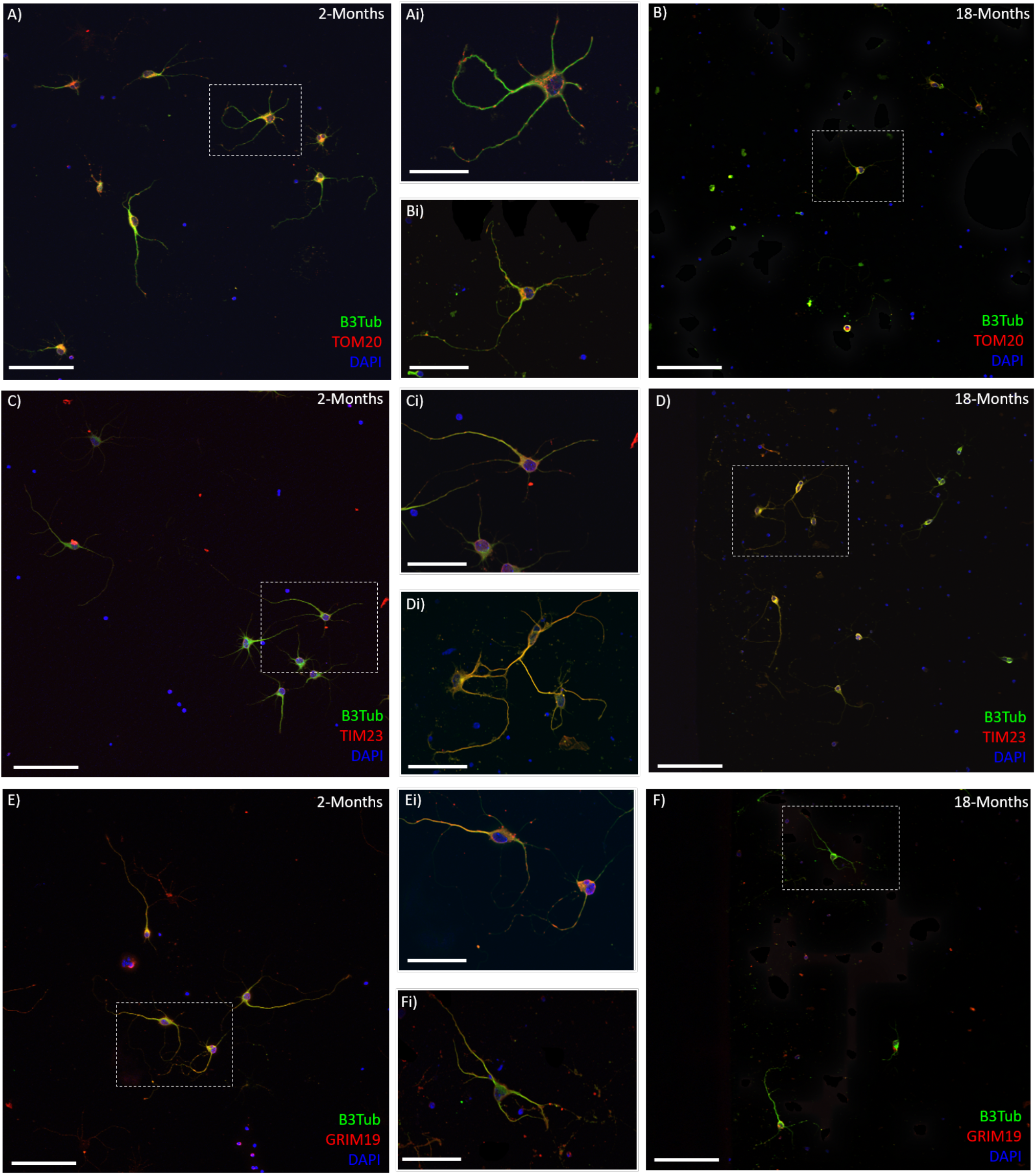
Mitochondrial membrane-associated proteins are altered in cortical neurons with age. A-F) Representative 20x confocol images of 2-month (A, C, E) and 18-month (B, D, F) cortical neurons after 7-days *in vitro;* stained with βIII-Tubulin (Green), DAPI (Blue), and either TOM20, TIM23 or GRIM19 (Red). Insets Ai-Fi) show higher magnification of single neurons or small clusters in young (Ai, Ci, Ei) and older (Bi, Di, Fi) cultures corrwsponding to the boxes on A-F. *Scale bars are 100μm for A-F and 50 μm for Ai-Fi*.

## 4. Discussion

The lack of functional recovery that is associated with spinal cord injury is linked to the degeneration of severed axons, the abnormal growth of what remains, and the failure of axons to regrow. Axonal growth inhibitors are also present after SCI, associated with elements of the secondary injury [6]. Consequently, axon growth has received a lot of attention in SCI research. Studies have investigated the mechanisms behind it, how it is affected by injury and, most significantly, how growth and regeneration can be promoted after injury. Despite rapid progress in understanding and manipulating the regulation of axon growth post-injury, we are yet to address how aging impacts axon growth and regeneration in the CNS. Mitochondria are vital to axon growth and are also known to decrease in efficacy and functionality with age, this poses a challenge where aging and SCI intersect.

This study represents the first comparison of cortical neurons from different aged brains, and a novel assessment of alterations in specific neuronal mitochondrial functions in these different ages. This data is the first to clearly demonstrate the age-dependent decline in cortical neurons neurite outgrowth *in vitro* and to suggest the neuron intrinsic properties are involved in this reduction. Indeed, neurite growth was reduced after 7-days *in vitro* using βIII-Tubulin as a marker. This does not allow for the identification of these neurites as either axons or dendrites and further specific staining is needed to explicitly demonstrate the age-dependent axon growth reduction in cortical neurons with this culture protocol. However, the observed decrease in neurite length and increase in branching in older cortical neurons is an indicator of the reduced growth potential of these cells, and may also lead to less efficient re-establishment of functional connections over the distance required in the case of axon damage seen in SCI. This work is also the first to specifically address the changes of mitochondrial functions with age in cortical neurons. Using this isolation system, we were able to culture cortical neurons from mice up to 33-months old (not shown). This will allow for a deeper understanding and characterization of the different neuron-intrinsic factors that are mediating the observed decline in growth and recovery potential seen in aging [7]. However, we acknowledge that these neurons are also impacted by a wide range of neuron-extrinsic factors [6] in the context of both normal aging and SCI, and that both neuron-intrinsic and extrinsic factors will have to be taken into account to fully comprehend the complexity of the different actors reducing axonal and dendritic growth in the CNS *in vivo*. Overall the mitochondrial changes presented here, even when showing trend without significance, support the hypothesis that mitochondrial functions in cortical neurons are reduced with age, which could, in part, explain the age-dependent decline in axon growth potential.

Mitochondrial DNA (mtDNA) is a multi-copy genome located in the mitochondrial matrix that encodes for 13 proteins of the ETC. An intact and functional mitochondrial genome is required for normal mitochondrial functioning. The damage and/or depletion of mtDNA, and the subsequent malfunction of mitochondria, has been implicated in a wide range of disorders including both aging and neurodegeneration [34]. A dysfunctional or damaged mtDNA genome have far reaching effects on the function and viability of mitochondria. Significantly; alterations, dysfunctions or decreases in mtDNA can have important downstream effects on the expression and function of the OXPHOS proteins of the ETC, and subsequent energy production. mtDNA copy number analysis compares select multi-copy mtDNA sequences to select multi-copy nuclear DNA sequences to act as an indicator of mitochondrial number in a cell/sample. In this paper, the multi-copy cytosolic ribosomal DNA sequence 18s and mitochondrial ribosomal DNA sequence 16s were used to estimate the ratio of mitochondrial to nuclear genomes, and therefore the number of mitochondrial to each cell. Surprisingly, this analysis showed increased relative expression of mtDNA in middle-aged and old mice compared to younger adults. This could have multiple implications for cortical neurons. Firstly, this increase could suggest a survival or compensatory mechanism for the older neurons. The increase in mtDNA may be an attempt to promote ETC protein expression and subsequently increase ATP production, and cell survival in response to the dysfunction state of neuronal mitochondria with age. Previous work in the lungs has shown an increase in mtDNA copy number in elderly individuals. This increase was linked to oxidative stress responses and suggested as a possible mechanism to compensate for defects in mitochondria retaining mutated mtDNA [35]. Other previous studies have consistently reported age-related decline in mtDNA copy number (mtDNAcn) [36–39] but are confounded by mixed cell samples and contamination from platelets, which inflate mtDNAcn and decline in number with age [40]. A study specifically examining peripheral blood mononuclear cells observed a positive correlation between mtDNAcn and age [40]. This may be used by cells as a compensatory upregulation to counteract the loss of functionally intact mitochondria with age [41, 42]. The increase in mtDNA expression, alongside the increased ATP levels, observed in isolated cortical motor neurons in the current study may be indicative of a similar compensatory mechanism to attempt to increase energy output from declining or dysfunctional mitochondria. This also fits with the maintenance of expression of OXPHOS complexes in older neurons as well, where we had expected a decrease. An increase in mtDNA has previously been shown to be beneficial in mitochondrial diseases. A previous study of heteroplasmic mtDNA mutations and disease found that an increase in wild type mtDNA ameliorated pathology across multiple tissues, while a reduction in mtDNA showed worsening pathology in postmitotic tissues but a beneficial effect in highly proliferative tissues [43]. In keeping with this idea, the cortex is a stable post-mitotic tissue and neurons do not proliferate, so the observed increase in mtDNA older neurons may be an attempt to rescue these cells and impede their decline. Secondly, this increase could indicate a reduction in mitophagy in older neurons that results in increased numbers of dysfunctional or declining mitochondria retained in the cells. The increase in TIM23 observed in older neurons may also support this idea. Another possibility is that the amount of mtDNA is linked to mitochondrial transport and the location of the mitochondria within the cell. During the dissociation process the cells’ axons and dendrites are sheared from the cell body, so the DNA extracted immediately from these isolated neurons will be largely that from the cell soma. The increase in mtDNA in older neurons may correlate with clustering of the mitochondria in the soma and decreased transport into the processes. In the current study TOM20 staining appeared to be less prominent in the neurites of older cortical neurons in culture, which may support this idea. Some previous papers have reported increased mitochondrial clustering around the nucleus when the cells are inclined to enter apoptosis [44–47]. This may also link to this hypothesis as aging cells are more stressed and vulnerable than their younger counterparts. Finally, this increase in mtDNA may be a cellular mechanism to compensate for a decline in expression, or dysfunction of, the protein machinery to synthesize OXPHOS proteins. These possibilities need to be further explored in older neurons.

The primary function of mitochondria is cellular respiration and energy production via the generation of ATP through OXPHOS. This process is significant in SCI for numerous reasons. Firstly, in the normal functioning CNS cortical neurons already rely on significant ATP production to maintain cellular functions [48, 49]. The formation of active axonal growth cones is required for neurons to promote axonal growth. This is essential for regeneration and restoration of any functionality after injury and requires even more substantial energy production [27]. Decreases in either the expression or function of OXPHOS complexes in the mitochondrial electron transport chain (ETC) in the aging CNS may have a significant impact on mitochondrial bioenergetics, and subsequently the ability to produce sufficient energy to support axon sprouting and growth. The observed age-related decline in mitochondrial activity [50–52] poses a challenge for SCI as successful axon regeneration requires an increase in mitochondrial dynamics [25, 26, 53], and the disruption of mitochondrial energy production is a significant detriment to axon growth. The loss of mitochondrial bioenergetics in aging may significantly compound SCI as this is also a characteristic of the injury, with significant decline in mitochondrial function observed as early as 12-hours after injury and progressively worsening [28].

Mitochondria in aging are considered to have decreased bioenergetics and declining function [10, 12, 54]. This led us to anticipate a decrease in the expression of some, if not all, of the five OXPHOS complexes in the ETC. Previously we have observed some decreases in Complex I and II, a small increase in Complex III in protein extracted from the entire cortex (not shown). However, in the enriched cortical neuron population examined in this study, we did not observe the anticipated decrease in expression of any of the five OXPHOS complexes with age. This could reflect that changes at the individual protein level are not occurring; however, it is possible that assembly of respiratory complexes or supercomplexes are impaired [55]. This was surprising as other measures examined suggested dysfunctional mitochondrial respiration. Using the Agilent Seahorse system to measure dynamic respiratory functions we observed reductions in basal respiration, maximal respiration, ATP production, and non-mitochondrial oxygen consumption in older neurons. We also observed an increase in non-mitochondrial oxygen consumption by older cortical neurons. These measures are indicative of the young neurons being healthier cells that are more able to produce the energy required for neurite growth in *in vitro* conditions. The older neurons, as well as having decreased respiratory measures, were unable to respond to the drugs used in the Mito Stress Test. This supports the idea that these cells are less metabolically active and also less able to respond to cellular stressors. The dysfunctional respiration in older neurons may be linked to the reduction in neurite growth, as well as the fragility of these cells in culture conditions. The extracellular acidification rate (ECAR) is associated with lactate efflux, an end-product of glycolysis, and points towards an active glycolytic metabolism. Increases in ECAR measurements indicate cellular activation and proliferation. The Seahorse showed an increased ECAR in younger neurons compared to their older counterparts, supporting a more energetic and active metabolism in these young cells. Using both the OCR and ECR measurements we observed a more quiescent metabolism in older neurons, compared to their more energetic younger counterparts. Taken together, the results from the OXPHOS western blot and the dynamic metabolic measurements from the Seahorse suggest that there is not a decrease in expression of OXPHOS Complexes in older cortical neuron mitochondria, but instead a dysfunction of respiration and energy metabolism in the existing ETC.

During OXPHOS incomplete coupling of mitochondria results in some dissipation of redox energy in a natural proton leak across the inner membrane. We also observed decreased proton leak in older cortical neurons. Previously, aging mitochondria from the rat liver, kidneys and heart have been seen to have decreased ΔΨM, and an increased rate of proton conductance [56]. Proton leak contributes to the resting metabolic rate of cells, with up to 20% of the basal metabolic rate being used to drive the basal leak [57], and decreased conductance can disturb energy homeostasis. Higher proton conductance can be linked to higher rates of OXPHOS, and mild uncoupling to protect against excessive ROS production [58]. The decreased proton leak in older neurons could be indicative of compromised energy homeostasis, be linked to the observed ΔΨM depolarization, and be another indication of dysfunctional respiration. Mitochondrial coupling efficiency is the proportion of oxygen consumed used to drive ATP synthesis compared with that driving the basal proton leak [59]. Decreases in coupling efficiency can be promoted by highly regulated uncoupling proteins to attenuate free radical production by mitochondria [57]. We observed a decrease in mitochondrial coupling efficiency in older neurons, which may be linked to the decreased proton leak and dysfunctional metabolism observed. This may also be associated with attempts to ameliorate the increased free radical production and accumulation seen in aging [13, 60].

The decline in energy production, energy expenditure, and basal metabolic rate with age has been documented for decades [61–63]. Following from this, the number of ATP molecules present in cortical neurons from each cohort is expected to decline with age, however, our results suggest that the number of ATP molecules per cortical neuron increases from young adult (2- and 6-month) to middle-aged (12-month) mice and remains elevated in aging. There may be several explanations for this surprising observation. First, the number of ATP molecules present is not directly connotative of the metabolic rate, expenditure, or efficiency of an organism Aging can reduce ATP/AMP ratio [64] suggesting reduced protein efficiency, concentration, or kinetics. Neurons in aged mice may have trouble utilizing free ATP, associated with decreases in their metabolic rate. Direct analysis of ATPase activity is required to elucidate age-related changes in neuronal energy metabolism. Secondly, it is possible that the elevated ATP levels recorded in cortical neurons with age, as well as the increased mtDNA expression, could be related to a compensatory upregulation of glycolysis [64], and perhaps mitochondrial biogenesis, to support the declining respiratory function. The lower membrane potential that was observed with age in these cortical neurons, as well as the reduced respiratory measurements and decreased ECAR seen in the Seahorse analysis are consistent with this idea. Thirdly, the assay used measures ATP present in the cell at the time of lysis rather than dynamic ATP production. Therefore, the absolute ATP numbers may be lower in younger mice due to a dysfunctional or slower metabolism of ATP in aging mice. This could indicate the increased storage but under-utilization of the ATP produced. Alternatively, for various reasons older animals may require increased ATP to perform compared to younger counterparts [65]. Finally, this observation may be linked to the euthanasia process for these mice before cell isolation. Mice are exposed to CO^2^ by inhalation for only as long as necessary for the mice to become unresponsive, prior to secondary physical euthanasia and the extraction of cortical tissue. The length of CO^2^ exposure may impact the amount of free ATP. The anaerobic environment of a CO^2^ chamber may induce rapid utilization of free ATP molecules in cortical neurons to prevent death. Hypoxia shifts ATP production towards glycolytic metabolism [66]. As aging increases glycolytic ATP production [64], the anaerobic environment of the CO^2^ chamber may be more favorable to aged neurons, and therefore, result in artificially higher ATP levels compared to younger counterparts. This data supports the contention that aging reduces the ability of cells and systems to react to changes in metabolic demand, and the ability to utilize stored energy. However, more in depth examination is required to elucidate the mechanisms underlying this surprising age-dependent increase, to clarify the mechanisms of ATP-storage and utilization in older cortical neurons and its biological implications for neuron growth and regeneration, especially in CNS trauma.

Changes in the mitochondrial membrane potential (ΔΨM) in older neurons may indicate an increased stress and dysfunctional state, which is likely to impair the ability of these mitochondria to efficiently produce the necessary energy and also contribute to increases in oxidative stress. ΔΨM is involved in mitochondrial respiratory rate, ATP synthesis, Ca^2+^ uptake and storage, and ROS balance, and is an important factor that can identify dysfunctional mitochondria [67, 68]. Mitochondria in aged neurons have Ca^2+^ buffering deficits as well as reduced ΔΨM [69, 70]. Alterations in Ca^2+^ homeostasis and decreased buffering capability are associated with both oxidative stress and neuronal excitotoxicity, a significant mechanism of axonal damage and an obstacle to axonal growth [14]. This study found decreased ΔΨM in middle-aged and older cortical neurons, compared to their younger adult counterparts. The decreased PE prevalence in the JC-1 stained neurons with age suggest increased membrane depolarization in older neurons. The JC-1 flow cytometry data in this study suggests that with age neurons have more dysfunctional mitochondria and are more sensitive to insult than younger neurons. This may have significant implications for the health and energy metabolism of these aging cells. ΔΨM is an important part of OXPHOS in that it is involved in energy storage in the mitochondria and, alongside the proton gradient, creates the transmembrane hydrogen ion gradient needed to produce ATP. Specifically, ΔΨM is important for transport of cations across the membrane in to the mitochondrial matrix and anions out [67]. The decreased ΔΨM seen in older cortical neurons, may be linked to the dysfunctional respiration and decreased ATP production that was also observed in these cells. Depolarization of ΔΨM is also linked to cellular apoptosis. Opening of the mPTP can induce ΔΨM depolarization, loss of oxidative phosphorylation, and release of apoptotic factors, and may be a precipitating event in apoptosis. Conversely, ΔΨM depolarization may be a consequence of the apoptotic-signaling pathway rather than an early precipitating factor [71]. In either instance, loss of ΔΨM can be considered an indicator of dysfunctional mitochondria and increased sensitivity to apoptotic insult. This is in keeping with the results of this study, and the suggested dysfunction and increased vulnerability of cortical neurons with age.

Membrane transport is vital to moving molecules into and out of the mitochondrial matrix and, as such, is linked to the ETC, the ΔΨM and proton gradient, as well ROS homeostasis [72, 73]. Two prominent families of multi-part membrane complexes are the translocase of outer membrane (TOM) [15, 16] and inner membrane (TIM) [17, 18]. TOM20 is a prominent component of the TOM complex responsible for the recognition and translocation of cytosolically synthesized mitochondrial preproteins. We observed a trend for decreased TOM20 expression in older cortical neurons. This may be an indicator of decreased or dysfunctional entry of proteins into the mitochondria. The TOM20 staining was clustered more prominently in the soma of the older neurons, with less expression in the processes. This may be an indicator of less mitochondria in the growing neurites and may also be a sign of declining cell health [47, 74]. Some previous studies have reported no change in TOM protein in aging skeletal muscle [75, 76], while other studies using mitochondrial fractionation have suggested an upregulation of TOM proteins in aging skeletal muscle [77]. The increase in TOM protein was particularly observed in older subjects that were functionally inactive [75], suggesting an upregulation in mitochondrial protein import may be a compensatory mechanism for other age-related dysfunction in skeletal muscle mitochondria with age. In this study, we observed a decrease in TOM20 expression with age in cortical neurons, however there was a significant increase in TIM23. This may be linked to a similar compensatory mechanism to try to boost dysfunctional aging mitochondria. While the TOM complex is the primary point of entry for most proteins destined for the mitochondrial matrix [78], TIM 23 is the primary translocase for matrix proteins to move through the inner mitochondrial membrane [17]. TIM23 is known to interact with subunits of the ETC and can play an important role in inserting inner-membrane proteins when the ΔΨM is reduced [47]. Over-expression of TIM23 in Parkinson’s Disease has been observed to reduce neurodegeneration. The neuroprotective mechanism of TIM23 increase may be linked to increased rate of protein import into mitochondria and also stabilization of complex I subunits [47]. Previously mitochondrial membrane transport, and especially TIM23, has been linked to neurodegenerative disorders, with data suggesting that deficient protein import is an early event in development of Huntington’s disease. This study observed that knocking down TIM23, using lentivirus-mediated delivery of shRNA to primary cortical and striatal neurons, resulted in increased neuronal death in a Huntington’s disease mouse model (HD transgenic R6/2 mice) [79]. In the current study we observed a significant increase in the expression of TIM23 in cortical neurons with age. This may indicate a compensatory mechanism for the dysfunctional aging mitochondria to help promote mitochondrial function and increase survival.

Gene associated with retinoid interferon induced cell mortality 19 (GRIM19) was first identified as an apoptotic nuclear protein associated with tumor cells [23]. To date, there have been multiple roles identified for GRIM19, including as a subunit to mitochondrial ETC Complex 1 [20, 23] and as a transport protein that helps to shuttle specific molecules to act in the mitochondria [19, 21, 80]. Significantly, in mitochondria, GRIM-19 is required for electron transfer activity of Complex I, and disruption of ΔΨM in GRIM-19 mutant models enhances sensitivity to apoptotic stimuli [22]. We observed a trend of decreased GRIM19 expression with age in cortical neurons. This decrease may be linked to the depolarized ΔΨM and dysfunctional respiration also observed. We observed no decrease in expression of OXPHOS Complex I with aging; however the decrease in GRIM19 may support a decline in function of the OXPHOS system rather than a decline in expression of the ETC proteins. The alteration in expression of these three proteins linked to mitochondrial membrane transport, ΔΨM and the ETC, suggest both another area of dysfunction in aging mitochondria as well as potential compensatory mechanisms for age-related decline.

Other elements of mitochondrial function in neurons have also been suggested to decline with age, potentially affecting axon growth and regeneration potential. Mitochondrial biogenesis, and the efficient turnover of appropriate proteins, is important for many elements of mitochondrial function including maintaining bioenergetics, ROS homeostasis, and fission/fusion. Dysfunctions in both mitochondrial biogenesis and in fission/fusion have been associated with the decline of mitochondria in aging [11, 81]. These dysfunctions may result in decreased mitochondrial functionality as well as decreased turnover of dysfunctional mitochondria, and could be caused by either changes in protein expression or in the maturation of the final products within the mitochondrial matrix [11]. Finally, the morphology of neurons presents unique challenges for energy homeostasis and distribution, especially projecting motor neurons with extremely long axons. Mitochondrial energy production is particularly in demand in neuronal synapses and at the growth cones of axons. This requires efficient mitochondrial trafficking and axonal transport to these specific, distal areas of the cell [82]. Axonal trafficking is essential for shuttling necessary supplies and energy to the growth cones and extending axons. This includes the transport of organelles (such as mitochondria), cytoskeletal proteins and other necessary components. Reduced speed of axon growth has been correlated with a diminished rate of axonal transport in the peripheral nervous system (PNS) with age. Enhancing mitochondrial trafficking may prove to be a promising target to increase bioenergetics and improve axon growth in an injury paradigm [83]. The age-related decline in mitochondrial transport, both in the PNS and the CNS, may have significant implications for the age-related decline in axon growth potential [7, 84]; further highlighting the importance of mitochondrial functioning and location in neurons after injury [83].

After SCI there are a wide range of injury cascades, mechanisms and dysfunctions exhibited; among these is mitochondrial dysfunction and oxidative stress [29]. Oxidative stress is a process that occurs when there is an imbalance between the production of ROS and the antioxidant systems required to detoxify these molecules. The mitochondria play a central role in monitoring and mediating this process in the cell. The significant role of oxidative stress and the formation of free radicals, such as reactive oxygen and nitrogen species, after SCI has been acknowledged for some time, and therefore examined in some detail [85–87]. A decline in mitochondrial activity and mitochondrial bioenergetics in the brain and spinal cord has been associated with normal aging, and has also been associated with a detrimental increase in ROS [88]. Oxidative stress and increased levels of free radicals, especially ROS, present in the injury environment after SCI inhibits axonal transport and promotes the degeneration of axons [14, 82, 89]. Increased ROS, particularly superoxide anions, has been linked to mtDNA damage and down-regulation of the ETC. This leads to subsequent mitochondrial dysfunctions and further promotes metabolic oxidative stress [90]. The CNS has a very active oxygen metabolism and high lipid content, the combination of which leave it particularly vulnerable to oxidative stress and free-radical mediated damages such as lipid peroxidation [85]. The disruption of the balance between OXPHOS and antioxidant systems, and oxidative stress are features of both the normal aging CNS as well as associated with injury and therefore can compound each other for a worse outcome for SCI patients.

Past studies have compared mitochondria isolated from brain tissue of animals to observe a plethora of age-related alterations [13]. Significantly, *In Vitro* studies using neurons [91] or astrocytes [92] from brains of both young and old mice, or cells “aged’’ in culture, have suggested that most cell types in the brain likely experience the accumulation of dysfunctional mitochondria during aging. This observation is borne out by the results of this current study that suggest an increase in mitochondrial number (as suggested by mtDNA expression) in aging, but dysfunctional indicators across the range of mitochondrial processes examined. It is likely that this decline in functional mitochondria, not a decline in mitochondrial number, is linked to the less efficient neurite growth observed here as well as the decreased growth potential after SCI in aging subjects. Moreover, it has been previously observed that in CNS injury the mitochondria translocate into injured axons resulting in an increase in the average mitochondria density [24]. A failure to increase mitochondrial function in neurons is linked to poor regeneration [24]. The retention of dysfunctional mitochondria seen in aging may also result in an increase in the number of mitochondria present, however this is not beneficial to the cell but rather may have a detrimental impact. Another possibility linked to decreased growth potential in older neurons is that the mitochondria in older neurons are not being transported to where they are needed to support axon growth. A recent study using a syntaphilin knock-out mouse model, lacking the static anchor protein to hold axonal mitochondria stationary and resulting in increased axonal mitochondrial motility, showed improvements in three distinct models of SCI and axonal injury [93]. This model suggested that enhancing mitochondrial transport and motility can enhance recovery after a SCI in young mice [93]. However, this contention has yet to be tested in an aging paradigm. The regulatory mechanisms that transport, distribute, and clear mitochondria in neurons are compromised in neurotrauma [94], which may compound the retention of dysfunctional mitochondria that is seen in aging. Mitochondrial energetics are beneficial to axon regeneration in SCI in young mice [93], however this has not been examined in middle-aged and aging animals. The differences in mitochondrial function and dynamics in older neurons may present a challenge, but also an opportunity to target to improve SCI recovery in aging patients.

Previously, mitochondria have been investigated as a potential therapeutic target in a range of disorders, most significantly in cancers. It is only in recent years that mitochondria-centric therapies, or ‘MitoCeuticals’ [95], have come to be explored for therapeutics in CNS injury. There are a number of reasons why targeting mitochondria therapeutically may be promising for the treatment of SCI [95]. Firstly, the secondary injury phase of SCI is a complex milieu involving many signaling cascades that intersect and diverge, and affect multiple targets [96, 97]. Due to the complex nature of this injury, targeting a single specific pathway or cascade may prove a less effective or insufficient approach to promote functional recovery after injury. This has led to the contention that a broader, sub-cellular approach may be more useful [98]. The therapeutic manipulation of mitochondria may be approached in a variety of ways, both genetically and pharmacologically, with a variety of outcomes [95, 98, 99]. Some examples of these that have begun to be explored include anti-apoptotic [100–102] as well as antioxidant strategies [103, 104], boosting mitochondrial bioenergetics [105] and biogenesis [99, 106], alternative biofuels [107–109], targeting the mPTP [30, 110–113], Ca2+ signaling and transmembrane channels. Of particular interest here, studies have even been conducted specifically to test pharmacological agents to target mitochondria in neurons [114]. In the case of SCI, ‘MitoCeuticals’ may prove to be an effective strategy to not only improve neuroprotection and create a more permissive lesion environment, but also potentially promote axon growth.

An interesting extension of the idea of using and manipulating mitochondria therapeutically in recent years has been the transplantation of whole mitochondria as a therapeutic agent [105, 115, 116]. Previous evidence in a variety of models, such as neurodegeneration and cardiac ischemia, suggests that transplantation of autologous mitochondria, either directly or systematically, is beneficial [116, 117]. The evolving idea that mitochondria that are released from cells after CNS injury can transfer to other cells and promote oxidative phosphorylation is the basis of using mitochondrial transplantation in SCI therapy [118, 119]. The theory behind this technique is that transplanted mitochondria will supplement existing antioxidant systems to encourage greater energy production and increased Ca^2+^ buffering efficiency [95]. These mitochondria-centric therapeutic strategies may be promising however, none of them have yet been explored in SCI in an aging paradigm.

## 5. Conclusions

Mitochondria are vital to the growth and regeneration of axons after injury, and this capability decreases in the aging CNS. The mitochondrial alterations we have observed in older neurons may play a significant role in the age-dependent decline in axon growth potential. In an increasingly older SCI population, this may have important implications as age and injury compound one another and result in worse injury outcomes for patients. The wide range of alterations, and the magnitude of such, does not point to one particular element but instead a general decline in function affecting aging neuronal mitochondria. This bears greater scrutiny in order to target mitochondria in aging neurons and pursue a potentially efficacious therapy for SCI in patients of all ages.

## Author Contributions

Conceptualization, TCS. and CGG.; methodology, TCS., A.S., D.H., Y.L., and T.H.; formal analysis, TCS.; data curation, TCS.; writing–original draft preparation, TCS.; writing–review and editing, TCS., A.S., T.H., Y.L., A.P.W., and CGG.; visualization, TCS.; supervision, CGG.; project administration, TCS.; funding acquisition, CGG. All authors have read and agreed to the published version of the manuscript.”

## Funding

This research was funded by a Craig H. Neilsen Foundation grant to Cedric Geoffroy (650332). Y.L. and A.P.W. were supported by NIH grant R01HL148153.

## Institutional Review Board Statement

This study was conducted according to the guidelines of the Declaration of Helsinki, and approved by the Institutional Review Board/Animal Ethics Committee of Texas A&M University (IACUC 2018-0324; 04/15/2019).

## Data Availability Statement

The data presented in this study are available on request from the corresponding author.

## Acknowledgments

We thank Amanda Manhke for assistance with the RT-qPCR.

## Conflicts of Interest

The authors declare no conflict of interest.

## Notes

### Competing Interest Statement

The authors have declared no competing interest.

